# Protected Polymorphisms and Evolutionary Stability of Patch-Selection Strategies in Stochastic Environments

**DOI:** 10.1101/003780

**Authors:** Steven N. Evans, Alexandru Hening, Sebastian J. Schreiber

## Abstract

We consider a population living in a patchy environment that varies stochastically in space and time. The population is composed of two morphs (that is, individuals of the same species with different genotypes). In terms of survival and reproductive success, the associated phenotypes differ only in their habitat selection strategies. We compute invasion rates corresponding to the rates at which the abundance of an initially rare morph increases in the presence of the other morph established at equilibrium. If both morphs have positive invasion rates when rare, then there is an equilibrium distribution such that the two morphs coexist; that is, there is a protected polymorphism for habitat selection. Alternatively, if one morph has a negative invasion rate when rare, then it is asymptotically displaced by the other morph under all initial conditions where both morphs are present. We refine the characterization of an evolutionary stable strategy for habitat selection from [Schreiber, 2012] in a mathematically rigorous manner. We provide a necessary and sufficient condition for the existence of an ESS that uses all patches and determine when using a single patch is an ESS. We also provide an explicit formula for the ESS when there are two habitat types. We show that adding environmental stochasticity results in an ESS that, when compared to the ESS for the corresponding model without stochasticity, spends less time in patches with larger carrying capacities and possibly makes use of sink patches, thereby practicing a spatial form of bet hedging.

## 1. Introduction

Habitat selection by individuals impacts key attributes of a population including its spatial distribution, temporal fluctuations in its abundance, and its genetic composition. In environmentally heterogeneous landscapes, individuals selecting more favorable habitats are more likely to survive or reproduce. As population densities increase in these habitats, individuals may benefit by selecting previously unused habitats. Thus, both environmental conditions and density-dependent feedbacks generate selective pressures on habitat selection. Under equilibrium conditions, spatial heterogeneity can select for populations exhibiting an ideal-free distribution–equal per-capita growth rates in all occupied patches and lower per-capita growth rates if individuals moved into unoccupied patches [Fretwell and Lucas, 1969]. Under non-equilibrium conditions, spatial-temporal heterogeneity can select for individuals occupying sink habitats in which the per-capita growth rate is always negative [Holt, 1997, Jansen and Yoshimura, 1998]. Environmental heterogeneity can also promote coexistence of genotypes only differing in their habitat choices [Jaenike and Holt, 1991]. Despite significant advances in the mathematical theory for habitat selection under equilibrium conditions, a mathematical theory for habitat selection in stochastic environments is largely lacking. Here, we take a step to addressing this mathematical shortfall while at the same gaining new insights into the evolution of habitat selection for populations living in stochastic, patchy environments.

Since the classic paper Fretwell and Lucas [1969], the ideal-free distribution has been studied extensively from empirical, theoretical, and mathematical perspectives. Empirical support for ideal-free distributions exists for many taxa including fish [Godin and Keenleyside, 1984, Oksanen et al., 1995, Haugen et al., 2006], birds [Harper, 1982, Doncaster et al., 1997], mammals [Beckmann and Berger, 2003], and insects [Dreisig, 1995]. For example, Oksanen et al. [1995] found that armored catfish in Panamanian stream pools were distributed such that the resource availability per catfish was equal in all occupied pools, despite significant variation in light availability across these occupied pools. Theoreticians have identified several “non-ideal” mechanisms (e.g. sedentarism, adaptive movement with finite speed, density-dependent dispersal) that, under equilibrium conditions, generate an ideal-free distribution [Hastings, 1983, Cosner, 2005, Gejji et al., 2012]. For example, at equilibrium, sedentary populations achieve an ideal-free distribution provided, paradoxically, the populations initially occupied all habitat patches. While many early studies asserted that the ideal free distribution is an evolutionarily stable strategy (ESS) [Fretwell and Lucas, 1969, van Baalen and Sabelis, 1993, Schreiber et al., 2000], only recent advanced nonlinear analyses fully verified this assertion [Cressman et al., 2004, Cressman and Křivan, 2006, 2010, Cantrell et al., 2007, 2010, 2012].

In nature, observed habitat occupancies are frequently less extreme than predicted by the ideal-free distribution: individuals underuse higher quality habitats and overuse lower quality habitats compared to theoretical predictions [Milinski, 1979, Tregenza, 1995]. Notably, populations occupying sink habitats have been documented in many species [Sokurenko et al., 2006, Tittler et al., 2006, Robinson et al., 2008, Anderson and Geber, 2010]. One possible explanation for these observations is that populations experience temporal as well as spatial variation in environmental conditions and, consequently, theory based on equilibrium assumptions tells an incomplete story. In support of this explanation, several theoretical studies have shown that occupation of sink habitats should evolve when temporal variation is sufficiently great in other habitats [Holt, 1997, Jansen and Yoshimura, 1998, Holt and Barfield, 2001, Schreiber, 2012]. These theoretical developments, however, rely on linearizations of density-dependent models, and do not analyze the dynamics of competing genotypes, the ultimate basis for evolutionary change due to natural selection. Hence, these studies leave unanswered the question, “Does the linear analysis correctly identify competitive exclusion in pairwise interactions that is the basis for the analysis of evolutionarily stable strategies?”

Within populations, individuals can exhibit different habitat selection strategies, and there is some evidence these differences can be genetically based [Via, 1990, Jaenike and Holt, 1991]. For instance, some individuals of the fruit fly species *Drosophila tripunctata* prefer tomato host plants (one potential habitat for its larvae) while others prefer mushrooms (another potential habitat), and these differences are based on two genetically independent traits, settling behavior and ovipositor site preference [Jaenike, 1985]. Jaenike and Holt [1991] found that genetic variation in habitat selection is common, especially in arthropods and mollusks. Furthermore, they demonstrated using mathematical models that this genetic variation can stem from density-dependent regulation occurring locally within each habitat. Specifically, Jaenike and Holt write “frequency-dependent selection favors alleles that confer upon their carriers a preference for underused habitats, even if there is no genetic variation in how well individuals are adapted to the different habitat” [Jaenike and Holt, 1991, p.S78]. Their analysis, however, doesn’t account for temporal fluctuations in environmental conditions and this raises the question, “Does environmental stochasticity facilitate or hinder the maintenance of genetic variation in habitat selection?”

To answer the aforementioned questions, we provide an in-depth analysis of a model introduced in [Schreiber, 2012]. The single genotype (i.e. monomorphic) version of this model and a characterization of its dynamics are given in Section 2. The competing genotype (i.e. dimorphic) version of the model and invasion rates of each genotype when rare are introduced in Section 3. In Section 4, we prove that these invasion rates determine the long-term fate of each of the genotypes. Specifically, if both genotypes have positive invasion rates when rare, then there is a positive stationary distribution under which the genotypes coexist. Alternatively, if one genotype has a negative invasion rate when rare, then it is asymptotically displaced by the other genotype. These result allows us to use the invasion rates when rare to explore conditions supporting a protected polymorphism for habitat selection. In Section 5, we refine the characterization of an evolutionary stable strategy for habitat selection from [Schreiber, 2012] in a mathematically rigorous manner, and provide an explicit formula for this ESS when there are two habitat types. Section 6 concludes with a discussion of how our results relate to the existing literature and identifies future challenges for the theory of habitat selection in stochastic environments.

## 2. The Monomorphic Model

To set the stage for two competing populations spread over several patches, we start with a single population living in one patch. Let *Z*_*t*_ be the population abundance at time *t* ≥ 0. The stochastic process (*Z*_*t*_)_*t*≥0_ is governed by the Itô stochastic logistic equation

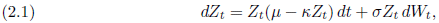

where *μ* is the intrinsic rate of growth of the population in the absence of stochasticity, *κ* is the strength of intraspecific competition, *σ*^2^ > 0 is the infinitesimal variance parameter of the stochastic growth rate, and (*W*_*t*_)_*t*≥0_ is a standard Brownian motion. The process (*Z*_*t*_)_*t*≥0_ is a strong Markov process with continuous paths. We call an object with such properties a *diffusion*.

As shown in our first proposition, the process (*Z*_*t*_)_*t*≥0_ lives in the positive half line ℝ_++_ := (0, ∞); that is, if we start it in a strictly positive state, then it never hits zero. Furthermore, the long-term behavior of the process is determined by the stochastic rate of growth 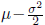. When the stochastic growth rate is negative the population abundance converges asymptotically to zero with probability one. On the other hand, when this parameter is positive the distribution of the abundance converges to an equilibrium given by a Gamma distribution. These results are well-known, but, as introduction to the methods used to prove our main results, we provide a proof in Appendix A.

### Proposition 2.1

*Consider the diffusion process* (*Z*_*t*_)_*t*≥0_ *given by the stochastic differential* *equation* (2.1).

- *The stochastic differential equation has a unique strong solution that is defined for all t* ≥ 0 *and is given by*

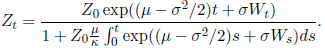
- *If Z*_0_ = *z* > 0, *then Z*_*t*_ > 0 *for all t* ≥ 0 *almost surely*.
- *If* 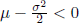*, then* lim_*t*→∞_ *Z*_*t*_ = 0 *almost surely*.
- *If* 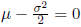*, then* lim inf_*t*→∞_ *Z*_*t*_ = 0 *almost surely*, lim sup_*t*→∞_ *Z*_*t*_ = ∞ *almost surely*, *and* 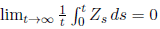 *almost surely*.
- *If* 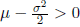*, then* (*Z*_*t*_)_*t*≥0_ *has a unique stationary distribution ρ on* ℝ_++_ *with Gamma density* 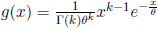*, where*

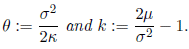 *Moreover, if Z*_0_ = *z* > 0, *then*

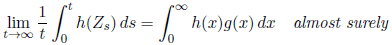

*for any Borel function h* : ℝ_++_ → ℝ *with* 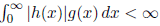. *In particular,*

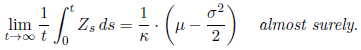

Next, we consider a population living in a spatially heterogeneous environment with *n* different patches. These patches may represent distinct habitats, patches of the same habitat type, or combinations thereof. The abundance of the population in the *i*-th patch at time *t* ≥ 0 is 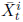. Let 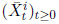 be given by

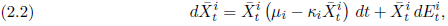

where *μ*_*i*_ is the intrinsic rate of growth the population in patch *i* in the absence of stochasticity, *κ*_*i*_ is the strength of intraspecific competition in patch *i*, and 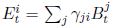 for a standard multivariate Brownian motion (*B*^1^, …, *B*^*n*^)^*T*^ on ℝ^*n*^ and an *n* × *n* matrix Г := (*γ*_*ij*_). The infinitesimal covariance matrix for the non-standard Brownian motion 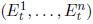 is Σ = (*σ*_*ij*_) := Γ^*T*^Γ.

The populations in the various patches described by equation (2.2) are coupled only by the spatial correlations present in the driving Brownian motion 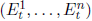. We further couple the population dynamics across patches by assuming the fraction of population in patch *i* eqauals *α*_*i*_ for all time. This spatial distribution can be realized at the scale of the individual when, as described in greater detail in Remark 2.2, individuals disperse rapidly and independently of one another in such a manner that the fraction of time spent in patch *i* equals *α*_*i*_ for each individual. Under this assumption, we call *α* = (*α*_1_, *α*_2_, …, *α*_*n*_) with *α*_*i*_ ≥ 0 for all 1 ≤ *i* ≤*n* and 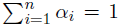, a **patch-selection strategy**. Continuing to denote the abundance of the population in the *i*-th patch at time *t* ≥ 0 as 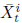, we have 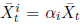, where 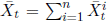 is the total population abundance at time *t* ≥ 0. If we impose these constraints on 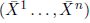, then it is heuristically reasonable that the process 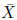 is an autonomous Markov process that satisfies the SDE

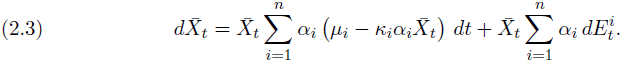

*Remark* 2.2. One way to justify the formulation of (2.3) rigorously is to first modify (2.2) to obtain a system of SDEs explicitly accounting for dispersal. Suppose that individuals disperse from patch *i* to patch *j* at a rate *δd*_*ij*_ for some fixed rate matrix *D* = (*d*_*ij*_). As usual, we adopt the convention *d*_*ii*_ = −∑_*j*≠*i*_*d*_*ij*_. The resulting system of SDEs is

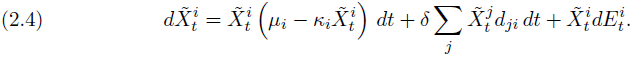

Assume that the rate matrix *D* has a unique stationary distribution *α*; that is, *α_j_* > 0 for 1 ≤ *j* ≤ *n* and 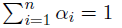,

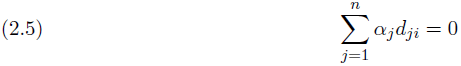

for 1 ≤ *i* ≤ *n*. In this case, a vector (*y*^1^, …, *y^n^*) satisfies

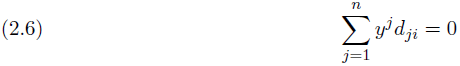

for 1 ≤ *i* ≤ *n* if and only if

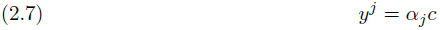

for 1 ≤ *j* ≤ *n* for some constant *c*. Moreover, summing (2.7) we find that

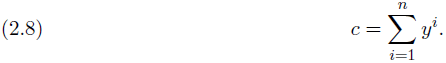

Note that by (2.5) we can write the drift term in (2.4) that contains *δ* as

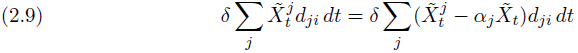

where 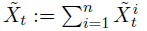. using (2.7) and (2.8), we see that (*x*^1^, …, *x*^n^) and 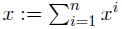 are such that

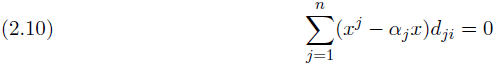

for *i* = 1, …, *n* if and only if

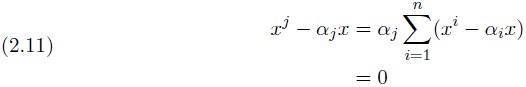

for 1 ≤ *j* ≤ *n*.

It follows from (2.9) and the equivalence between (2.10) and (2.11) that as *δ* increases the solution of (2.4) experiences an increasingly strong drift towards the one-dimensional subspace

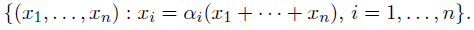

In the limit *δ* → ∞, it is plausible that the system (2.4) converges to one for which

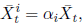

where 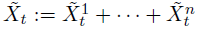, and the total population size 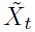 satisfies the autonomous one-dimensionl SDE (2.3) with 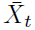 replaced by 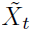. This heuristic for obtaining (2.3) as a high dispersal rate limit of (2.4) can be made rigorous by applying Theorem 6.1 from Katzenberger [1991].

Let 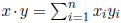 denote the standard Euclidean inner product and define another inner product 〈·, ·〉_*κ*_ by 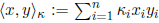. Since (*α* · *E*_*t*_)_*t*≥0_ is a Brownian motion with infinitesimal variance parameter *α* · Σ*α*, (2.3) can be expressed more simply as

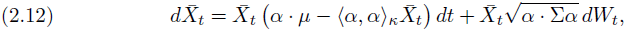

where *W*_*t*_ is a standard Brownian motion.

The total population 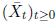 defined by (2.12) behaves **exactly** like the one-patch case defined by (2.1) with the parameters *μ* → *μ* · *α*, *κ* → 〈*α*, *α*〉_*κ*_ and 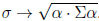. In particular, 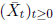 is a diffusion process and we have the following immediate consequence of Proposition 2.1

### Proposition 2.3

*Consider the diffusion process* 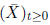 *given by* (2.12).

- *If* 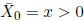*, then* 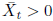 *for all t* ≥ 0 *almost surely*.
- *If* 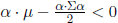*, then* 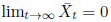 *almost surely*.
- *If* 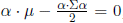*, then* 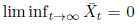 *almost surely,* 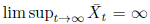 *almost surely, and* 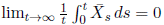 *almost surely*.
- *If* 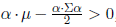*, then the process* 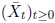 *has a unique stationary distribution* 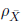 *on* ℝ_++_ *with Gamma density* 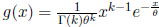*, where*

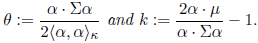 *Moreover*,

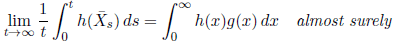

*for any Borel function h* : ℝ_++_ → ℝ *with* 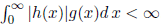. *In particular,*

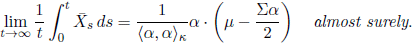

For the dynamics (2.2) in patch *i*, Proposition 2.1 implies that if there was no coupling between patches by dispersal, then then population abundance in patch *i* would converge to 0 if *μ*_*i*_ − *σ*_*ii*_/2 < 0 and converge to a non-trivial equilibrium if *μ*_*i*_ − *σ*_*ii*_/2 > 0. As noted by Schreiber [2012], and illustrated below, the spatially coupled model is such that the population can persist and converge to an equilibrium even when *μ*_*i*_ − *σ*_*ii*_/2 < 0 for all patches.

**Persistence of coupled sink populations in symmetric landscapes.** Consider a highly symmetric landscape where *μ*_*i*_ = *r*, *σ*_*ii*_ = σ^2^ > 0 for all *i*, *κ*_*i*_ = *a* for all *i*, and *σ*_*ij*_ = 0 for all *i* ≠ *j*. If individuals are equally distributed across the landscape (*α*_*i*_ = 1/*n* for all *i*), then

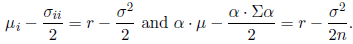

The increase in the stochastic growth rate from *r* − *σ*^2^/2 for an isolated population to *r* − *σ*^2^/(2n) for the spatially coupled population stems from individuals spending equal time in patches with uncorrelated environmental fluctuations. Specifically, the environmental variance experienced by individuals distributing their time equally amongst *n* uncorrelated patches is *n* times smaller than the environmental variance experienced by an individual spending their time entirely in one patch. Whenever *σ*^2^ > 2*r* > *σ*^2^/*n*, this reduction in variance allows the entire population to persist despite patches, in and of themselves, not supporting population growth.

## 3. Dimorphic model and invasion rates

To understand the evolution of patch-selection strategies, we now consider competition between populations that only differ in their patch-selection strategy. Let *X*_*t*_ and *Y*_*t*_ be the total population sizes at time *t* ≥ 0 of two populations playing the respective patch selection strategies *α* = (*α*_1_, *α*_2_, …, *α*_*n*_) and *β* = (*β*_1_, *β*_2_, …, *β*_*n*_), so that the densities of the populations in patch *i* are *α*_*i*_*X*_*t*_ and *β*_*i*_*Y*_*t*_ at time *t* ≥ 0. The dynamics of these two strategies are described by the pair of stochastic differential equations

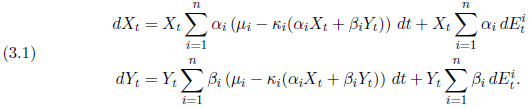

Since

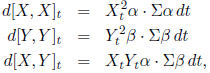

the diffusion process ((*X*_*t*_, *Y*_*t*_))_*t*≥0_ for the spatially coupled, competing strategies can be represented more compactly as

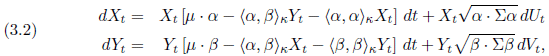

where (*U*, *V*) is a (non-standard) Brownian motion with covariance structure *d*[*U*, *U*]_*t*_ = *dt*, *d*[*V*, *V*]_*t*_ = *dt*, and 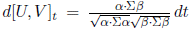. Using a construction similar from Remark 2.2, system (3.2) can be seen as a high dispersal limit. This system exhibits a degeneracy when *U* = *V* i.e. 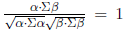. If Σ is nonsingular, then, by the Cauchy-Schwarz inequality, this degeneracy only occurs if *α* = *β*. We do not consider this possibility in what follows.

To determine whether the two populations coexist or one displaces the other, we introduce the invasion rate *ℐ*(*α*, *β*) of a population playing strategy *β* when introduced at small densities into a resident population playing strategy *α*. As shown in the next proposition, this invasion rate is defined by linearizing the dynamics of *Y* and computing the long-term population growth rate *ℐ*(*α*, *β*) associated with this linearization. When *ℐ*(*α*, *β*) > 0, the population playing strategy *β* tends to increase when rare. When *ℐ*(*α*, *β*) < 0, the population playing strategy *β* tends to decrease when rare.

### Proposition 3.1

*Consider the partially linearized system*

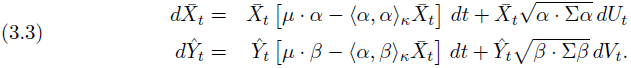

Assume 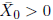 and *Ŷ*_0_ > 0.

*If α* · (*μ* − Σ*α*/2) > 0, *so the Markov process* 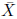 *has a stationary distribution concentrated on* ℝ_++_*, then the limit* 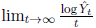 *exists almost surely and is given by*

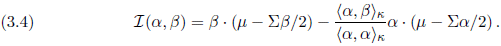

*On the other hand, if α* · (*μ* − Σ*α*/2) ≤ 0, *so that* 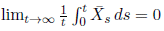 *almost surely, then the limit* 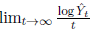 *exists almost surely and is given by*

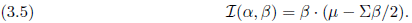

*Proof.* By Itô’s lemma,

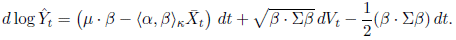

Assume that 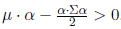. By Proposition 2.3,

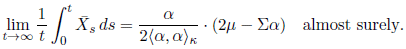

Therefore,

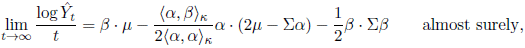

as claimed.

On the other hand, assume that 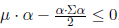. By Proposition 2.3,

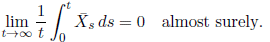

Therefore,

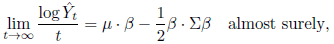

again as claimed.

In the next proposition, we show that if a population playing strategy *β* cannot invade a population playing strategy *α* (i.e. *ℐ*(*α*, *β*) < 0), then the population strategy *α* can invade the population playing strategy *β* (i.e. *ℐ*(*β*, *α*) > 0). This suggests, as we will show in the next section, that such a strategy *α* should exclude strategy *β*.

### Proposition 3.2

*Suppose that α* · (*μ* − Σ*α*/2) > 0 *and ℐ*(*α*, *β*) < 0. *Then, ℐ*(*β*, *α*) > 0.

*Proof*. Set *A* := *α* · (*μ* − Σ*α*/2) and *B* := *β* · (*μ* − Σ*β*/2). Assume that *A* > 0 and *ℐ*(*α*, *β*) < 0. To show that *ℐ*(*β*, *α*) > 0, we consider two cases, *B* ≤ 0 and *B* > 0. Suppose *B* ≤ 0. Then, *ℐ*(*β*, *α*) = *A* > 0 by Proposition 3.1 and by assumption.

Alternatively, suppose that *B* > 0. Then

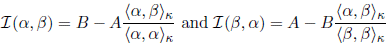

by Proposition 3.1. Assume, contrary to our claim, that *ℐ*(*α*, *β*) < 0 and *ℐ*(*β*, *α*) ≤ 0. From the Cauchy-Schwarz inequality 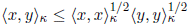 we get

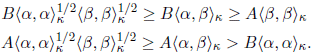

The above inequalities yield the contradiction 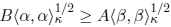 and 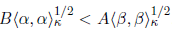.

An immediate consequence of Proposition 3.1 is the following corollary. This corollary implies that if a population playing strategy *β* can invade a population playing strategy *α* and a population playing strategy *α* can invade a population playing strategy *β*, then a single population playing strategy *α* converges to a non-trivial equilibrium and the same is true of a single population playing strategy *β*. This suggests, as we show in the next section, that under these conditions these two strategies should coexist.

### Corollary 3.3

*The invasion rate satisfies ℐ*(*α*, *β*) ≤ *β* · (*μ* − Σ*β*/2). *In particular, if ℐ*(*α*, *β*) > 0 *and ℐ*(*β*, *α*) > 0, *then α* · (*μ* − Σ*α*/2) > 0 *and β* · (*μ* − Σ*β*/2) > 0.

## 4. Exclusion and protected polymorphisms

Our main results about the dimorphic process (*X*, *Y*) is that the invasion rates determine the long-term fate of competing strategies. If the invasion rates predict that strategy *β* cannot invade a population playing strategy *α*, then the population playing strategy *α* drives the population playing strategy *β* asymptotically to extinction as shown in the following theorem. We give a proof in Appendix B.

### Theorem 4.1

*If α* · (*μ* − Σ*α*/2) > 0 *and ℐ*(*α*, *β*) < 0, *then, for* 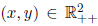, *the probability measures*

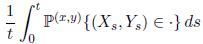

*converge weakly as t* → ∞ *to* 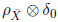*, where* 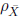 *is the unique stationary distribution of* 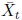 *concentrated on* ℝ_++_, *and δ*_0_ *is the point mass at* 0.

On the other hand, if the invasion rates predict that each strategy can invade when rare, then the following theorem proves that the competing strategies coexist: for any initial conditions the joint distribution of (*X_t_*, *Y_t_*) converges as *t* → ∞ to a probability distribution on 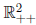 with density *ψ* and, moreover, for any Borel set 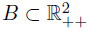 the long term proportion of times *t* for which (*X_t_*, *Y_t_*) spends in B converges to

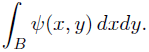

A proof is given in Appendix C. In order to appreciate the assumptions of the theorem, it helps to recall Corollary 3.3 which says that if ℐ(*α*, *β*) > 0 and ℐ(*β*, *α*) > 0 then *α*·(*μ* − Σ*α*/2) > 0 then *β*·(*μ* − Σ*β*/2) > 0 so that a single population playing strategy *α* or *β* will persist.

### Theorem 4.2

*Suppose that ℐ*(*α*, *β*) > 0 *and ℐ*(*β*, *α*) > 0. *Then, there exists a unique stationary distribution π of* (*X*, *Y*) *on* 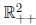 *that is absolutely continuous with respect to Lebesgue measure. Moreover, for any bounded, measurable function* 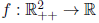

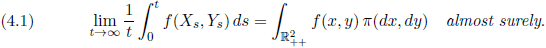

*Furthermore, the process* (*X*, *Y*) *is strongly ergodic, so that for any initial distribution m one has*

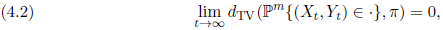

*where d*_TV_ *is the total variation distance.*

From the perspective of population genetics, the coexistence of these two strategies corresponds to a **protected polymorphism:** each strategy (a morph) increases when rare and, therefore, is protected from extinction. This protection from extinction, however, is only ensured over ecological time scales as mutations may result in new morphs that can displace one or both coexisting morphs [Ravigné et al., 2004]. The concept of protected polymorphisms was introduced by Prout [1968] when studying deterministic models of competing haploid populations in a spatially heterogenous with overlapping generations. Turelli et al. [2001] extended this concept to stochastic difference equations for competing haploid populations with a constant population size. Theorem 4.2 provides a mathematically rigorous characterization of protected polymorphisms for our stochastic models with fluctuating population sizes.

Theorem 4.2 implies that coexistence depends on the intrinsic stochastic growth rate of the populations and the competitive effect of each population on the other. The intrinsic stochastic growth rates are given by

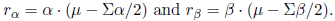

While the competitive effect of the population with strategy *α* on the population with strategy *β* is given by the ratio of the magnitude of *α* projected in the *β* direction (i.e. 〈*β*/∥*β*∥_*κ*_, *α*〉_*κ*_ where 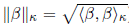) divided by the magnitude of *β* (i.e. ∥*β*∥_*κ*_). Mathematically, the competitive effect of *α* on *β* and the competitive effect of *β* on *α* are given by

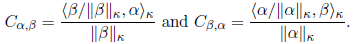

Provided *r*_*α*_ and *r*_*β*_ are positive, Theorem 4.2 implies that there is a protected polymorphism if

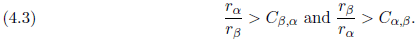

In words, the relative intrinsic stochastic growth rate of each population must exceed the competitive effect on itself due to the other population. Conversely, if one of the inequalities in (4.3) is reversed, then Theorem 4.1 implies that one population excludes the other. Unlike the standard Lotka-Volterra competition equations, Proposition 3.2 implies that both inequalities in (4.3) cannot be simultaneously reversed and, consequently, bistable dynamics are impossible.

**Environmental stochasticity impedes protected polymorphisms in symmetric landscapes.** Consider a landscape where all patches have the same carrying capacities (e.g. *κ*_*i*_ = 1 for all *i*), the same intrinsic rates of growth (i.e. *μ*_*i*_ = *a* for all *i*), and the same amount of uncorrelated environmental stochasticity (e.g. *σ*_*ii*_ = *σ*^2^ for all *i* and *σ*_*ij*_ = 0 for *i* ≠ *j*). Then the protected polymorphism inequalities (4.3) become

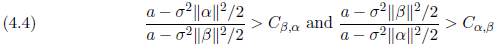

where the only *σ*^2^ dependency is on the left hand sides of both inequalities. As 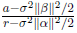 is a decreasing function of *σ*^2^ whenever 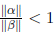 and an increasing function of *σ*^2^ whenever 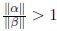, it follows that the set of set of strategies supporting a protected polymorphism

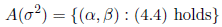

is a decreasing function of *σ*^2^ i.e 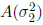 is a proper subset of 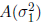 whenever *σ*_2_ > *σ*_1_ ≥ 0. Figure 1A illustrates this conclusion for a two-patch landscape. Intuitively, increasing environmental stochasticity in these symmetric landscapes reduces the stochastic growth rate for all strategies and, thereby, makes it less likely for populations to persist let alone coexist. For asymmetric landscapes, how the set *A*(*σ*^2^) of protected polymorphisms varies with *σ*^2^ is more subtle, as illustrated in Figure 1B. In this case, some protected polymorphisms are facilitated by environmental stochasticity, while other protected polymorphisms are disrupted by environmental stochasticity.

**Figure 1.**
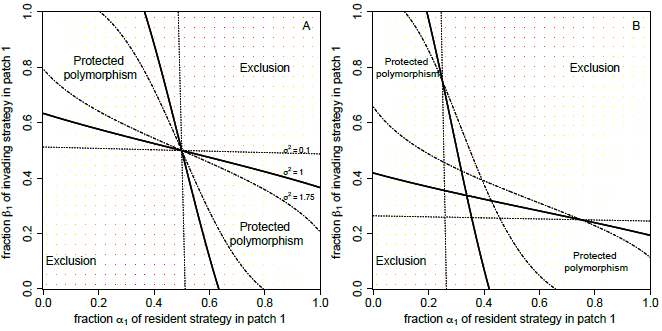
Protected polymorphisms and exclusion in two-patch landscapes. Contour plots of *ℐ*(*α*, *β*) where lighter shades correspond to higher values of *ℐ*(*α*, *β*). The regions where *ℐ*(*α*, *β*)*ℐ*(*β*, *α*) > 0 are delineated by the solid curves and correspond to parameter combinations supporting a protected polymorphism. Regions where *ℐ*(*α*, *β*)*ℐ*(*α*, *β*) < 0 correspond to strategies that cannot coexist. The dashed-dotted and dotted
curves indicate how regions of coexistence and exclusion change for higher and lower levels of environmental stochasticity *σ*^2^, respectively. In panel A, the landscape is spatially homogeneous with *μ* = (1, 1), *κ* = (1, 1) and Σ = *σ*^2^*I* where *I* is the 2 × 2 identity matrix. In B, the landscape is spatial heterogeneous with respect to the deterministic carrying capacities *κ* = (3, 1) and the remaining parameters as A.

For the symmetric landscapes, we can identify a strategy that displaces all others. Namely, the strategy 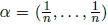 of visiting all patches with equal frequency. This strategy maximizes the function function *α* ↦ *a* − *σ*^2^∥*α*∥^2^/2. Hence, if we consider a competing strategy (*β* ≠ *α*, then 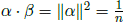 and

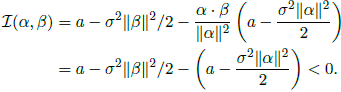

e.g. the invasion rates are negative along the vertical transect *α*_1_ = 1/2 in Figure. 1A. This strategy *α* is an example of an evolutionarily stable strategy that we discuss further in the next section.

## 5. Evolutionarily stable strategies

The concept of an evolutionary stable strategy was introduced by Maynard Smith and Price [1973]. Loosely stated, an evolutionary strategy is a strategy that cannot be invaded by any other strategy and, consequently, can be viewed as an evolutionary endpoint. For our models, we say patch selection strategy *α* is an *evolutionarily stable strategy (ESS)* if *ℐ*(*α*, *β*) < 0 for all strategies *β* ≠ *α*. In light of Theorem 4.1, an ESS not only resists invasion attempts by all other strategies, but can displace all other strategies. An ESS *α* is called a *pure ESS* if *α*_*i*_ = 1 for some patch *i*, otherwise it is a *mixed ESS*. Our next result provides an algebraic characterization of mixed and pure ESSs. However, it remains to be understood whether these ESSs can be reached by small mutational steps in the strategy space (i.e. are convergently stable [Geritz et al., 1997]).

### Theorem 5.1

*Assume that the covariance matrix* Σ *is positive definite and that there is at least one patch selection strategy which persists in the absence of competition with another strategy; that is, that* max_*α*_ *α* · (*μ* − Σ*α*/2) > 0.

**Mixed strategy:** *An ESS α with α*_*i*_ > 0 *for i* ∈ *I with I* ⊆ {1, 2, …, *n*} *and* ∣*I*∣ ≥ 2 *satisfies*

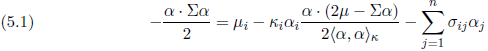

*for all i* ∈ *I*. *Conversely, if* ∣*I*∣ = *n, then a strategy α satisfying* (5.1) *is an ESS*.

**Pure strategy:** *The strategy α*_*i*_ = 1 *and α*_*j*_ = 0 *for j* ≠ *i is an ESS if and only if*

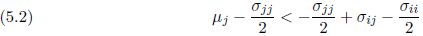

*for all j* ≠ *i*.

*Furthermore, in the case of n* = 2*, there exists a mixed ESS whenever the reversed inequalities*

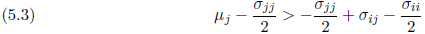

*hold for i* = 1, 2 *and j* ≠ *i*.

The first statement of Theorem 5.1 provides a sufficient and necessary condition for a mixed ESS utilizing all patches. For example, in a symmetric landscape (as described in the previous section), this ESS condition is only satisfied for *α* = (1/*n*, 1/*n*, …, 1/*n*).

The second statement of Theorem 5.1 provides a characterization of when using only a single patch is an ESS. Since the right hand side of equation (5.2) is negative, using patch *i* can only be an ESS if all other patches have a negative stochastic rate of growth, 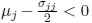 for all *j* ≠ *i*. However, even if only patch *i* has a positive stochastic growth rate, an ESS may use the other patches, as we illustrate next for two-patch landscapes.

### ESSs in two-patch, uncorrelated landscapes

For an uncorrelated two patches landscape (i.e. *n* = 2 and *σ*_12_ = 0), Theorem 5.1 implies that there is a mixed ESS whenever

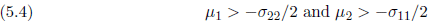

and this ESS satisfies

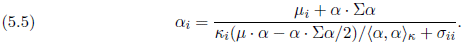

Equation (5.4) implies that even if deterministic growth in patch 2 is strictly negative (i.e. *μ*_2_ < 0), then there is selection for movement into this patch provided the variance of the fluctuations in patch 1 are sufficiently large relative to the intrinsic rate of decline in patch 2 (Fig. 2).

**Figure 2.**
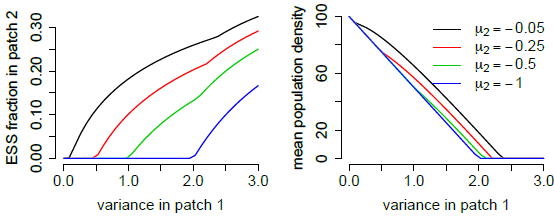
ESS for patch selection (left) and mean population abundance (right) in a source-sink landscape. Parameter values: *n* = 2, *σ*_11_ = *σ*^2^, *σ*_22_ = *σ*_12_ = 0, *μ* = (1, *μ*_2_), and *κ* = (1, 1).

In the limit of no noise (i.e. *σ*_*ii*_ ↓ 0 for *i* = 1, 2), equation (5.5) becomes

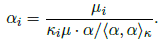

While our results do not apply to the deterministic case, this limiting expression for the ESS suggests, correctly, that the ESS for the deterministic model satisfies

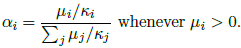

In other words, the fraction of individuals selecting patch *i* is proportional to the equilibrium density *μ*_*i*_/*κ*_*i*_ supported by patch *i*. Equation (5.5) implies that adding stochasticity in equal amounts to all patches (i.e. *σ*_*ii*_ = *σ*^2^ for all *i*) results in an ESS where, relative to the deterministic ESS, fewer individuals select patches supporting the highest mean population abundance and more individuals selecting patches supporting lower mean population abundances (Fig. 3).

**Figure 3.**
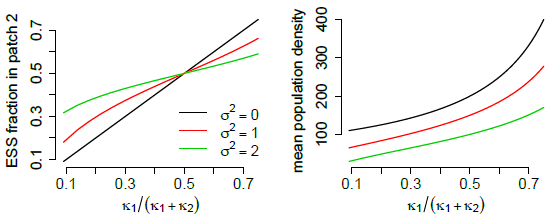
The effect of the deterministic carrying capacities and environmental stochasticity on the ESS for patch selection (left) and mean population abundance (right) in a two-patch landscape. The ratio *κ*_1_/(*κ*_2_ + *κ*_2_) corresponds to the ratio of the deterministic carrying capacity (*μ*_2_/*κ*_2_) in patch 2 to the sum of the deterministic carrying capacities (*μ*_1_/*κ*_2_ + *μ*_2_/*κ*_2_) when *μ*_1_ = *μ*_2_ = 1. Parameter values: *n* = 2, *σ*_11_ = *σ*_22_ = *σ*^2^, *σ*_12_ = 0, *μ* = (1, 1), and *κ* = (1, *κ*_2_).

## 6. Discussion

Habitat selection by organisms is a complex process determined by a mixture of genetic, developmental, ecological, and environmental factors. For ecologists, habitat selection plays a fundamental role in determining the spatial and temporal distribution of a population [Rosenzweig, 1981, Orians and Wittenberger, 1991]. For evolutionary biologists, habitat selection determines the suite of environmental factors driving local adaptation [Edelaar and Bolnick, 2012]. Indeed, in the words of the eminent evolutionary biologist Ernst Mayr, “With habitat and food selection – behavioral phenomena – playing a major role in the shift into new adaptive zones, the importance of behavior in initiating new evolutionary events is self-evident” [Mayr, 1963, p. 604]. Here, we examined how spatial and temporal heterogeneity in demographic rates across multiple habitat patches influence the dynamics of competing populations who only differ in their habitat patch selection preferences. We assume that habitat selection has a genetic basis (e.g. genes that influence the physiological or neurological capacity of individuals to detect and respond to habitat cues) and that genetic differences in habitat choice have no pleiotropic effects on habitat specific fitness. Our analysis reveals that, generically, only two outcomes are possible, coexistence or displacement of one population by the other for all initial conditions, and that these outcomes are determined by the invasion rates of populations when rare. In addition to providing a mathematically rigorous justification of prior work, our analysis provides new insights into protected polymorphisms for habitat selection and raises several questions about evolutionary stable strategies for habitat selection.

Protected polymorphisms correspond to populations of competing genotypes exhibiting negative frequency-dependence: each population tends to increase when rare [Prout, 1968]. In the case of patch selection, these competing populations differ in the frequencies in which they select habitat patches. In a survey of the empirical literature, Jaenike and Holt [1991] found “that genetic variation for habitat selection is common, especially in arthropods and mollusks, the groups that have been studied most frequently.” Moreover, they argued that some of this variation may be maintained through protected polymorphism. Specifically, “in a haploid model without intrinsic fitness differences among genotypes [i.e. soft selection], genetic variation in fixed habitat preferences may be maintained stably” [Jaenike and Holt, 1991, pg. S83]. We provide a general analytic criterion (see, inequality (4.3)) characterizing these protected polymorphisms for spatially and temporally variable environments. This criterion depends on the intrinsic fitnesses (*r*_*α*_ and *r*_*β*_) of each population and their competitive coefficients (*C*_*α*,*β*_ and *C*_*β*,*α*_) that characterize the effect of each population on the other. Competitive effects are greatest when there is an overlap in patch use and one population tends to select the patches with the higher carrying capacities more than the other population. Intuitively, by occupying patches with a larger carrying capacities, populations achieve higher regional densities. Coupled with overlap in patch use, these higher densities result in a greater competitive impact of one population on another. A protected polymorphism occurs when the relative fitness of each population (e.g. *r*_*α*_/*r*_*β*_ for strategy *α*) is greater than the competitive effect of the other population on it (e.g. *r*_*α*_/*r*_*β*_ > *C*_*β*,*α*_ for the population playing strategy *α*). Hence, as in the case of species coexistence [Chesson, 2000], protected polymorphism are most likely when fitness differences are small (i.e. *r*_*α*_/*r*_*β*_ ≈ 1) and competitive effects are small (i.e. both *C*_*α*,*β*_ and *C*_*β*,*α*_ < 1). Environmental stochasticity solely effects the intrinsic fitness terms and can facilitate or inhibit protected polymorphisms. For landscapes in which all patches experience the same degree of uncorrelated, temporal variation, environmental stochasticity has an inhibitory effect as it magnifies fitness differences between competing strategies (e.g. *r*_*α*_/*r*_*β*_ increases with environmental stochasticity). For asymmetric landscapes, however, temporal variability can facilitate polymorphisms by reducing fitness differences of competing strategies.

In contrast to protected polymorphisms, our analysis reveals that populations playing an evolutionarily stable strategy (ESS) for patch selection not only thwart invasion attempts by all other strategies but also can invade and displace a population playing any other strategy. Furthermore, our analysis provides a mathematically rigorous justification of an earlier characterization of ESSs [Schreiber, 2012]. This characterization implies that populations playing the ESS always occupy source habitats (i.e. patches where 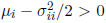). Indeed, consider a population playing strategy *α* that does not occupy some source patch, say patch *i*. Then a different behavioral genotype *β* that only selects patch *i* can invade as 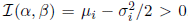. In the limiting case of a deterministic environment, our characterization of the ESS recovers the classic result of McPeek and Holt [1992]: the fraction of time spent in a patch is proportional to the carrying capacity of the patch. Adding environmental stochasticity generally results in populations playing the ESS decreasing the time spent in the patches with larger carrying capacities and possibly making use of sink patches (i.e. patches where 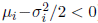). This shift in patch choice can be viewed as a spatial form of bet hedging: individuals increase fitness by decreasing the variance in their stochastic growth rate at the expense of their mean growth rate [Childs et al., 2010].

We are able to show that for two patch landscapes there always exists an ESS for patch selection. However, several questions remain unanswered. First, what happens for landscapes with more than two patches? Is there always an ESS? Second, while we know that a population playing an ESS can displace a monomoprhic population playing a different strategy, can it displace polymorphic populations? Finally, are ESSs always convergently stable [Geritz et al., 1997]? If there are positive answers to this final suite of questions, then ESSs can be generally viewed as the ultimate evolutionary end state for patch selection strategies.

Going beyond the models considered here, studying the evolution habitat use faces many challenges. Our models assume that populations spend a fixed fraction of time in each patch and do so instantaneously. What happens if we relax these assumptions? For example, if populations are more ideal and able to track changes in population density instantaneously, then we have something closer to the classical notion of ideal free movement [Fretwell and Lucas, 1969]. For these populations, what is the optimal (in an evolutionary sense) density-dependent strategy? Moreover, can such a strategy displace the static strategies considered here? Alternatively, if populations are less ideal and diffusing randomly on the landscape, what happens then? The linear version of this question was tackled in part by Evans et al. [2013]. However, the mathematical analysis for analogous stochastic models with density-dependent feedbacks is largely unexplored. Going beyond single species, the coevolution of patch selection among interacting species has a rich history for spatially heterogeneous, but temporally homogeneous environments [van Baalen and Sabelis, 1993, Křivan, 1997, Schreiber et al., 2000, van Baalen et al., 2001, Schreiber et al., 2002, Cressman et al., 2004, Schreiber and Vejdani, 2006, Cantrell et al., 2007]. For example, spatial heterogeneity can select for the evolution of contrary choices in which the prey prefers low quality patches to escape the predator and the predator prefers high quality patches to capture higher quality food items [Fox and Eisenbach, 1992, Schreiber et al., 2000]. Understanding how environmental stochastic influences this coevolution of patch choice and the community level consequences of these coevolutionary outcomes provides a plethora of important, yet largely untouched challenges for future work.

## Appendix A Proof of Proposition 2.1

The stochastic differential equation for *Z* is of the form

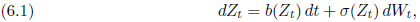

where *b*(*z*) := *μz* − *κz*^2^ and *σ*(*z*) := *σz*. It follows from Itô’s existence and uniqueness theorem for strong solutions of stochastic differential equations that this equation has a unique strong solution up to possibly a finite but strictly positive explosion time.

Set *R*_*t*_ := log *Z*_*t*_ for *t* ≥ 0. By Itô’s lemma,

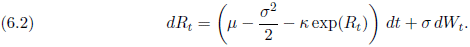

It follows from the comparison principle of Ikeda and Watanabe (see Chapter VI Theorem 1.1 of Ikeda and Watanabe [1989]), Theorem 1.4 of Le Gall [1983], or Theorem V.43.1 of Rogers and Williams [2000]) that

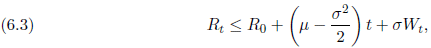

and so *Z* does not explode to +∞ in finite time. Moreover, since *r* ↦ *μ* − *κe*^*r*^ is a bounded, uniformly Lipschitz function on (−∞, 0] it follows from Itô’s existence and uniqueness theorem that *R* does not explode to −∞ in finite time, so that *Z* does not hit 0 in finite time. We could have also established this result by using the scale function and speed measure calculated below to check Feller’s necessary and sufficient for the boundary point of a one-dimensional diffusion to be inaccessible − see Theorem 23.12 of Kallenberg [2002].

It is not hard to check using Itô’s lemma that an explicit solution of the SDE is

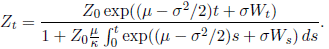

We see from the inequality (6.3) that if *μ* − *σ*^2^/2 < 0, then lim_*t*→∞_ *Z*_*t*_ = 0 almost surely.

We use the theory based on the scale function and speed measure of a one-dimensional diffusion (see, for example, Chapter 23 of Kallenberg [2002] or Sections V.6-7 of Rogers and Williams [2000]) below to establish that *Z* is positive recurrent with a unique stationary distribution when *μ* − *σ*^2^/2 > 0. Similar calculations show that *Z* is null recurrent when *μ* − *σ*^2^/2 = 0, and hence lim inf_*t*→∞_ *Z*_*t*_ = 0 almost surely and lim sup_*t*→∞_ *Z*_*t*_ = ∞. It follows from (6.2) and the comparison principle that if *Z′* and *Z″* are two solutions of (6.1) with respective parameters *μ′*, *κ′*, *σ′* and *μ″*, *κ″*, *σ″* satisfying *μ′* ≤ *μ″*, *κ′* = *κ″*, *σ′* = *σ″* and the same initial conditions, then 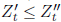. We will show below that

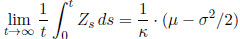

almost surely when *μ* − *σ*^2^/2 > 0, and hence

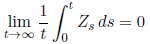

almost surely when *μ* − *σ*^2^/2 = 0.

We now identify the scale function and speed measure of the one-dimensional diffusion *Z*. A choice for the scale function is

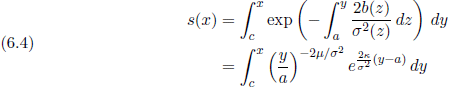

for arbitrary *a*, *c* ∈ ℝ_++_ (recall that the scale function is only defined up to affine transformations). If we set 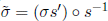, then

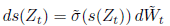

and the diffusion process *s*(*Z*) is in natural scale on the state space *s*(ℝ_++_) with speed measure *m* that has density 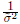.

The total mass of the speed measure is

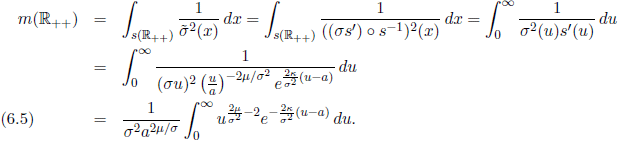

By Theorem 23.15 of Kallenberg [2002], the diffusion process *Z* has a stationary distribution concentrated on ℝ_++_ if and only if the process *s*(*Z*) has (−∞, +∞) as its state space and the speed measure has finite total mass or *s*(*Z*) has a finite interval as its state space and the boundaries are reflecting. The introduction of an extra negative drift to geometric Brownian motion cannot make zero a reflecting boundary, so we are interested in conditions under which *s*(ℝ_++_) = (−∞, ∞) and the speed measure has finite total mass. We see from (6.4) and (6.5) that this happens if and only if *μ* − *σ*^2^/2 > 0, a condition we assume holds for the remainder of the proof.

The diffusion *s*(*Z*) has a stationary distribution with density 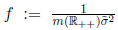 on *s*(ℝ_++_) = (−∞, +∞), and so the stationary distribution of *Z* is the distribution on ℝ_++_ that has density

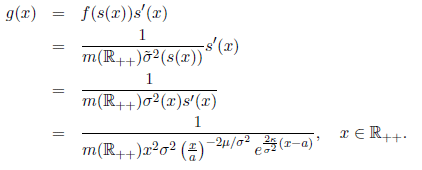

This has the form of a Gamma(*κ*, *θ*) density with parameters 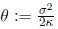 and 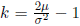. Therefore,

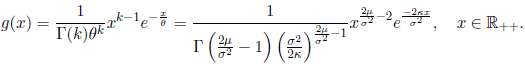

Theorem 20.21 from Kallenberg [2002] implies that the shift-invariant *σ*-field is trivial for all starting points. The ergodic theorem for stationary stochastic processes then tells us that, if we start *Z* with its stationary distribution,

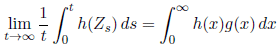

for any Borel function *h* : ℝ_++_ → ℝ with 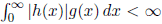. Since *Z* has positive continuous transition densities we can conclude that

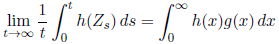

ℙ^*x*^-almost surely for any *x* ∈ ℝ_++_.

In particular,

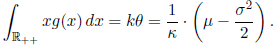

## Appendix B Proof of Theorem 4.1

To simplify our presentation, we re-write the joint dynamics of *X* and *Y* as

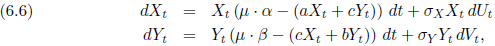

where 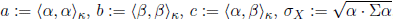, and 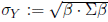.

To prove Theorem 4.1, we need several preliminary results. First, we prove existence and uniqueness of solutions to the system (6.6) as well as a useful comparison result in Theorem 6.1. Second, in Proposition 6.3, we establish that (*X*_*t*_, *Y*_*t*_) remains in 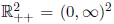 for all *t* ≥ 0 whenever 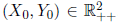. Third, in Proposition 6.4, we show that weak limit points of the empirical measures 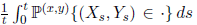 are stationary distributions for the process (*X*, *Y*) thought of as a process on 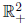 (rather than 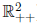). Finally, we show that lim_*t*→∞_ *Y*_*t*_ = 0 with probability one in Proposition 6.5 and conclude by showing that 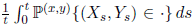 converges weakly to 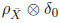 concentrated on ℝ_++_ × {0}.

### Theorem 6.1

*The stochastic differential equation in* (6.6) *has a unique strong solution and X*_*t*_*, Y*_*t*_ ∈ *L*^*p*^ (ℙ^(*x*,*y*)^) *for all t, p* > 0 *for all* 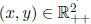. *This solution satisfies X*_*t*_ > 0 *and Y*_*t*_ > 0 *for all t* ≥ 0, ℙ^(*x*,*y*)^-*almost surely for all* 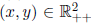. *Let* 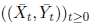 be the stochastic process defined by the pair of stochastic differential equations

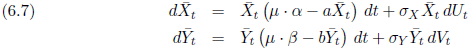

*If* 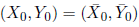*, then*

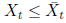

*and*

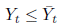

*for all t* ≥ 0.

*Proof.* The uniqueness and existence of strong solutions is fairly standard, see, for example, Theorem 2.1 in Li and Mao [2009]. One notes that the drift coefficients are locally Lipschitz so strong solutions exist and are unique up to the explosion time. It is easy to show this explosion time is almost surely infinite (see Theorem 2.1 in Li and Mao [2009]). Next, suppose that 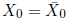. We adapt the comparison principle of Ikeda and Watanabe (Chapter VI Theorem 1.1 from Ikeda and Watanabe [1989]) proved by the local time techniques of Le Gall (see Theorem 1.4 from Le Gall [1983] and Theorem V.43.1 in Rogers and Williams [2000]) to show that 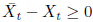 for all *t* ≥ 0.

Define *ρ* : ℝ_+_ → ℝ_+_ by *ρ*(*x*) = |*x*|^2^. Note that

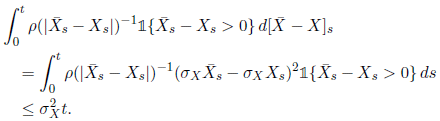

Since ∫_0+_*ρ*(*u*)^−1^ *du* = ∞, by Proposition V.39.3 from Rogers and Williams [2000] the local time at 0 of 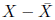 is zero for all *t* ≥ 0. Put *x*^+^ := *x* ∨ 0. By Tanaka’s formula (see equation IV.43.6 in Rogers and Williams [2000]),

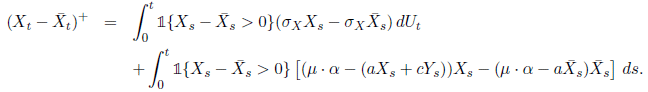

For *K* > 0 define the stopping time

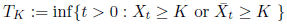

and the stopped processes 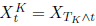 and 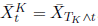. Then, stopping the processes at *T*_*K*_ and taking expectations yields

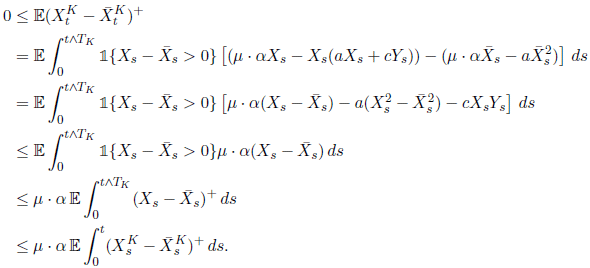

By Gronwall’s Lemma (see, for example, Appendix 5 of Ethier and Kurtz [2005]) 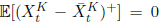 for all *t* ≥ 0, so 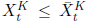 for all *t* ≥ 0. Now let *K* → ∞ and recall that 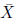 does not explode to get that 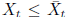 for all *t* ≥ 0. Since we have shown before that 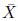 is dominated by a geometric Brownian motion, a process that has finite moments of all orders, we get that *X*_*t*_, *Y*_*t*_ ∈ *L*^*p*^(ℙ^(*x*,*y*)^) for all *t*, *p* > 0 and for all 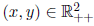.

*Remark* 6.2. Note that the SDEs for all the processes considered here have unique strong solutions in *L*^*p*^ for all *t* ≥ 0, *p* > 0 and for all strictly positive starting points. This follows by arguments similar to those that are in Theorem 2.1 from Li and Mao [2009] and in Theorem 6.1 by noting that our SDEs for (*X*, *Y*), 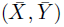 etc. are all of the form

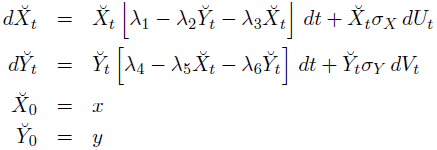

for *λ*_1_, …, *λ*_6_ ∈ ℝ_+_ and *x*, *y* ∈ ℝ_++_

The next proposition tells us that none of our processes hit zero in finite time.

### Proposition 6.3

*Let* (*X*, *Y*) *be the process given by* (6.6). *If* 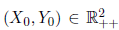, *then* 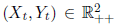 *for all t* ≥ 0 *almost surely. A similar conclusion holds for all of the other processes we work with*.

*Proof.* As an example of the method of proof, we look at the process (*X*, *Y*) given by (6.6). Taking logarithms and using Itô’s lemma,

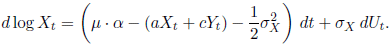

Therefore,

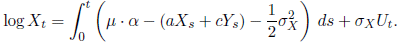

can’t go to −∞ in finite time because *X*_*t*_ and *Y*_*t*_ do not blow up.

### Proposition 6.4

*Let* (*X*, *Y*) *be the process given by* (6.6) *and fix* 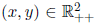. *Any sequence* {*t*_*n*_}_*n*∈*ℕ*_ *such that t*_*n*_ → ∞ *has a subsequence* {*u*_*n*_}_*n*∈ℕ_ *such that the sequence of probability measures*

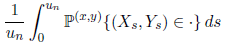

*converges in the topology of weak convergence of probability measures on* 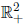. *Any such limit is a stationary distribution for the process* (*X*, *Y*) *thought of as a process with state space* 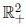.

*Proof.* Set *φ*(*x*, *y*) := *x* + *y* so that *φ* ≥ 0 for *x*, *y* > 0. Put *ψ*(*x*, *y*) = *μ* · *αx* + *μ* · *βy* − *x*(*ax* + *cy*) − *y*(*cx* + *by*). Note that *ψ* is bounded above on the quadrant *x*, *y* ≥ 0 and lim_∥(*x*, *y*)∥→∞_ *ψ*(*x*, *y*) = −∞ where ∥·∥ is the Euclidean distance on ℝ^2^. Using Itô’s lemma we get

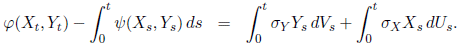

Therefore, 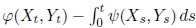 is a martingale. Applying Theorem 9.9 of Ethier and Kurtz [2005] completes the proof.

The following result is essentially Theorem 10 in Liu et al. [2011]. We include the proof for completeness.

### Proposition 6.5

*Suppose that α*·*μ*−*α*·Σ*α*/2 > 0, *β*·*μ*−*β*·Σ*β*/2 > 0, *and ℐ*(*α*, *β*) < 0. *If* (*X*, *Y*) *is the process given by* (6.6)*, then* lim_*t*→∞_ *Y*_*t*_ = 0 ℙ^(*x*,*y*)^-*a.s. for all* 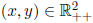.

*Proof*. Using Ito’s lemma and the definition of *ℐ*(*α*, *β*),

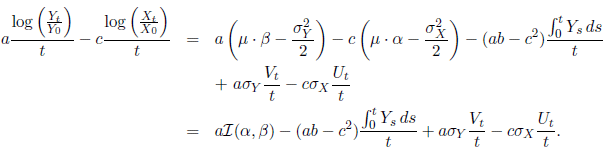

By the Cauchy-Schwarz inequality, (*ab* − *c*^2^) = 〈*α*, *α*〉_*κ*_〈*β*, *β*〉_*κ*_ − (〈*α*, *β*〉_*κ*_)^2^ ≥ 0, and so

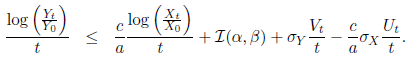

Let 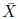 be the process defined by (6.7) with 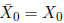. Proposition 2.3 implies

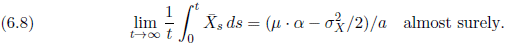

It follows from Theorem 6.1 that 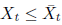 for all *t* ≥ 0. Thus, with probability one,

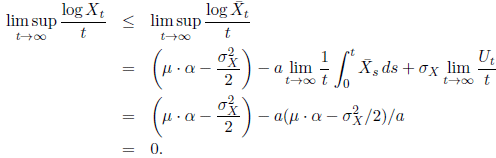

Since *U* and *V* are Brownian motions, 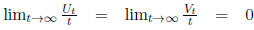, and 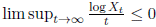 almost surely, so

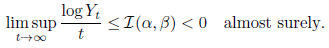

In particular, lim_*t*→∞_ *Y*_*t*_ = 0 almost surely.

We can now finish the proof of Theorem 4.1. Fix *ε* > 0 and *η* > 0 sufficiently small. Define the stopping time

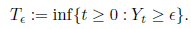

and the stopped process 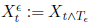. By Proposition 6.5, there exists *T* > 0 such that

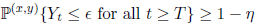

Define the process 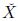 via

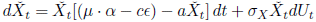

and the stopped process 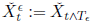. Start the process 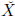 at time *T* with the condition 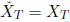. We want to show that the process 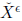 is dominated by the process *X*^*ε*^, that is 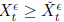 for all *t* ≥ *T*. By the strong Markov property, we can assume *T* = 0.

The proof is very similar to the one from Theorem 6.1. With the notation from the proof of Theorem 6.1, we have

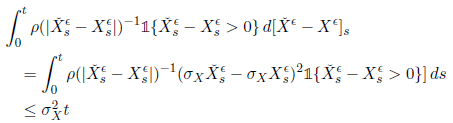

so the local time of the process 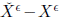 at zero is identically zero. Then, using Tanaka’s formula,

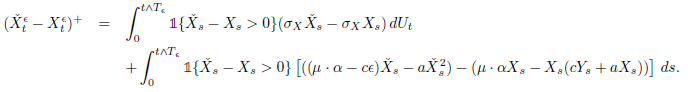

Taking expectations,

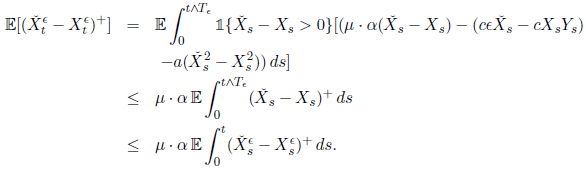

By Gronwall’s Lemma, 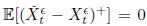. As a result, remembering we assumed *T* = 0, we have 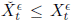 for all *t* ≥ *T*. For *ε* small enough we know that 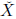 has a stationary distribution concentrated on ℝ_++_. For any sequence *a*_*n*_ → ∞, if the Cesaro averages 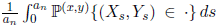 converge weakly, then the limit is a distribution of the form *φ* ⊕ *δ*_0_, where *φ* is a mixture of the unique stationary distribution 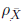 described in Proposition 2.3 and the point mass at 0. By the above, the limit of 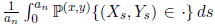 cannot have any mass at (0, 0) because 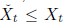 on the event {*Y*_*t*_ ≤ *ε* for all *t* ≥ *T*} that has probability ℙ^(*x*,*y*)^{*Y*_*t*_ ≤ *ε* for all *t* ≥ *T*} ≥ 1 − *η*. Since *η* > 0 was arbitrary, we conclude that 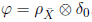, as required.

## Appendix C Proof of Theorem 4.2

Our proof is along the same lines as the proofs of Theorems 4 and 5 in Schreiber et al. [2011].

We will once again simplify our notation by re-writing the SDE for the pair (*X*, *Y*) as in (6.6). We assume throughout this appendix that the hypotheses of Theorem 4.2 hold; that is, *ℐ*(*α*, *β*) > 0 and *ℐ*(*β*, *α*) > 0.

Let 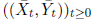 be the stochastic process defined by the pair of stochastic differential equations in (6.7) with initial conditions 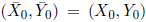. We know from Theorem 6.1 that 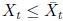 and *Y*_*t*_ ≤ Ȳ_*t*_ for all *t* ≥ 0.

Note from Corollary 3.3 that *α* · (*μ* − Σ*α*/2) > 0 and *β* · (*μ* − Σ*β*/2) > 0 and hence, by Proposition 2.3, the process 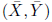 has a unique stationary distribution on 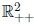 and is strongly ergodic.

Let

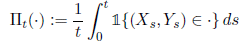

be the normalized occupation measures of (*X*, *Y*). We know that the random probability measures

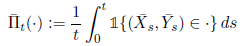

converge almost surely and so, in particular, they are tight on 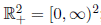; that is, for any *ε* > 0 we can find a box [0, *K*] × [0, *K*] such that

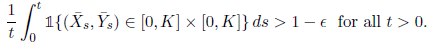

Therefore,

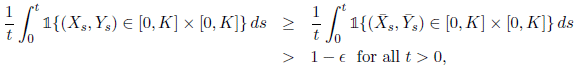

and hence the normalized occupation measures of (*X*, *Y*) are also tight on 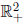. By Prohorov’s theorem [Kallenberg, 2002, Theorem 16.3], there exists a random probability measure *ν* on 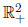 and a (possibly random) sequence (*t*_*n*_) ⊂ ℝ_++_ such that *t*_*n*_ → ∞ for which

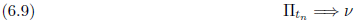

as *n* → ∞ almost surely, where ⇒ denotes weak convergence of probability measures on 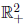. That is, with probability one for all bounded and continuous function *u*: 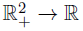 we have

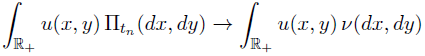

as *n* → ∞.

### Proposition 6.6

*The probability measure ν is almost surely a stationary distribution for* (*X*, *Y*) *thought of as a process with state space* 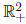.

*Proof.* Let (*P*_*t*_)_*t*≥0_ be the semigroup of the process (*X*, *Y*) thought of as a process on 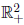. For simplicity let us write *Z*_*t*_ := (*X*_*t*_, *Y*_*t*_) for all *t* ≥ 0 and 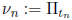.

By the Strong Law of Large Numbers for martingales, we have that for all *r* ∈ ℝ_+_ and all bounded measurable functions *f*

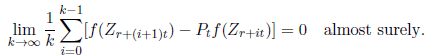

As a result,

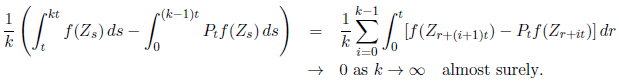

This implies that

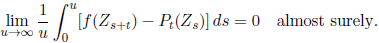

Thus,

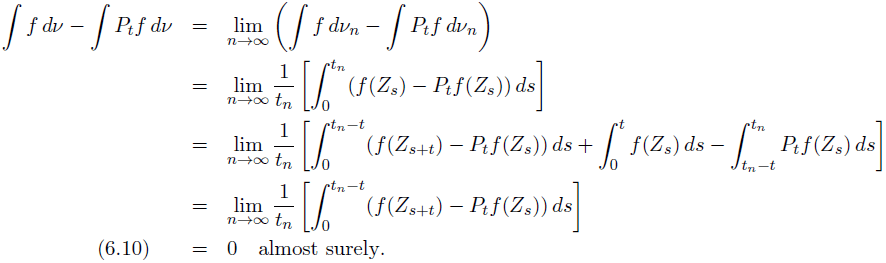

The last result is equivalent to saying that *ν* is almost surely a stationary distribution for (*X*, *Y*).

### Proposition 6.7

*There exists a stationary distribution π of* (*X*, *Y*) *that assigns all of its mass to* 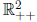.

*Proof.* We argue by contradiction. Because the process stays in one of the four sets ℝ_++_, ℝ_++_ × {0}, {0} × ℝ_++_, {(0, 0)} when it is started in the set, any stationary distribution for (*X*, *Y*) thought of as a process on 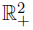 can be written as a convex combination of stationary distributions that respectively assign all of their masses to one of the four sets, should such a stationary distribution exist for the given set. Suppose there is no stationary distribution that is concentrated on 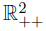. Then, any stationary distribution is the convex combination of stationary distributions that respectively assign all of their mass to the three sets ℝ_++_ × {0}, {0} × ℝ_++_, and {(0, 0)}, and hence any stationary distribution is of the form

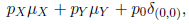

where the random variables *p*_*X*_, *p*_*Y*_, *p*_0_ are nonnegative and *p*_*X*_ + *p*_*Y*_ + *p*_*0*_ = 1 almost surely, and 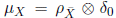 and *μ*_*Y*_ = *δ*_0_ ⊗ *ρ*_*Ȳ*_ for 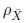 and *ρ*_*Ȳ*_ the unique stationary distributions of 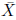 and *ρ*_*Ȳ*_. Next, we proceed as in Proposition 6.5 to find the limit of 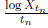. Let us first argue that

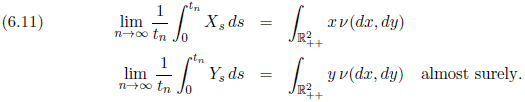

Note that the infinitesimal generator of (log *X*, log *Y*) thought of as a process on ℝ^2^ is uniformly elliptic with smooth coefficients and so it has smooth transition densities (see, for example, Section 3.3.4 of Stroock [2008]). Moreover, an application of a suitable minimum principle for the Kolmogorov forward equation (see, for example, Theorem 5 in Section 2 of Chapter 2 of Friedman [1964]) shows that the transition densities are everywhere strictly positive. It follows that (*X*, *Y*) thought of as a process on 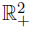 has smooth transition densities that are everywhere positive.

Because the process 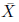 also has smooth, every positive transition densities for similar reasons, the almost sure behavior of the 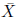 started from a fixed point is the same as it is starting from its stationary distribution 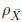. As a result, we get by Birkhoff’s pointwise ergodic theorem [Kallenberg, 2002, Theorem 10.6] that, for all *K* > 0,

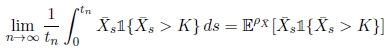

ℙ^*x*^ almost surely for any *x* ∈ ℝ_+_. Therefore, by dominated convergence

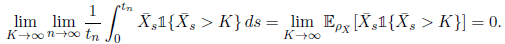

The following inequalities are immediate due to the positivity of the terms

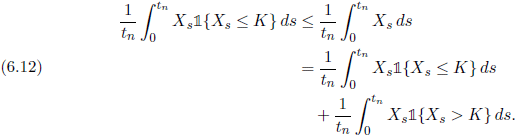

Recall that 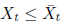 for all *t* ≥ 0 and hence

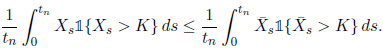

This implies

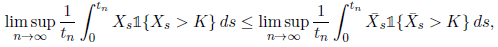

and therefore

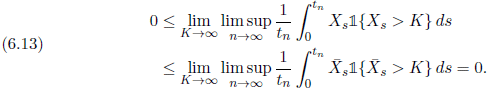

By (6.9), and Theorem 4.27 of Kallenberg [2002],

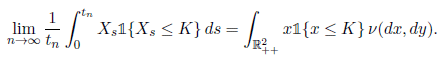

for any *K* such that

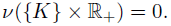

While this last condition need not hold a priori for all *K*, we can only have

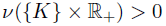

for countably many *K*, so there exists a sequence (*K_m_*) ⊂ ℝ_+_ such that *K_m_* → ∞ as *m* → ∞ with

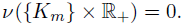

By dominated convergence,

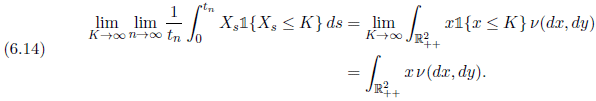

Combining (6.12), (6.13) and (6.14) gives (6.11).

It follows from Itô’s formula, the observation *ℐ*(*α*, *α*) = 0, (6.11), and the fact that 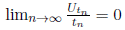 that

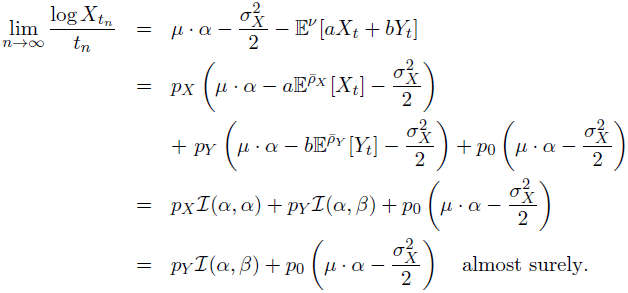

By assumption, *ℐ*(*α*, *β*) > 0 and we have already observed that 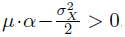. Because 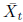 converges in distribution as *t* → ∞ to a distribution that assigns all of its mass to 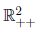, it follows that 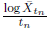 converges in probability to 0. However, since 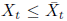 for all *t* ≥ 0 it follows that 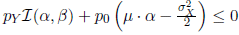 and hence

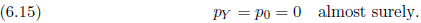

The same argument applied to (*Y*_*t*_)_*t*≥0_ establishes

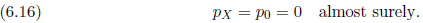

Therefore, *p*_*X*_ = *p*_*Y*_ = *p*_0_ = 0, and this contradicts the assumption that *p*_*X*_ + *p*_*Y*_ + *p*_0_ = 1.

We can now finish the proof of Theorem 4.2.

*Proof.* Proposition 6.7 implies that (*X*, *Y*) has a stationary distribution π on 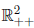. By Theorem 20.17 from Kallenberg [2002], our process (*X*, *Y*) is either Harris recurrent or uniformly transient. We say that (*X*_*t*_, *Y*_*t*_) → ∞ almost surely as *t* → ∞ if 1_*K*_(*X*_*t*_, *Y*_*t*_) → 0 as *t* → ∞ for any compact set 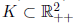. Theorem 20.21 from Kallenberg [2002] gives that if (*X*, *Y*) is transient, then (*X*_*t*_, *Y*_*t*_) → ∞ and so (*X*, *Y*) cannot have a stationary distribution. Hence, since we know our process has a stationary distribution *π*, it must be Harris recurrent. Theorem 20.21 from Kallenberg [2002] then gives us equation (4.1).

Theorem 20.18 from Kallenberg [2002], 20.18 gives that any Harris recurrent Feller process on 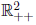 with strictly positive transition densities has a locally finite invariant measure that is equivalent to Lebesgue measure and is unique up to a normalization. We already know that we have a stationary distribution, so this distribution is unique and has an almost everywhere strictly positive density with respect to Lebesgue measure. Theorem 20.12 from Kallenberg [2002] says that any Harris recurrent Feller process is strongly ergodic, and so equation (4.2) holds.

*Remark* 6.8. In Theorem 3.1 of Zhang and Chen [2013], the authors claim to show that the system of SDE describing (*X*, *Y*) always has a unique stationary distribution. We note that their use of moments just checks tightness in 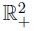 := [0, ∞)^2^ and not in 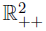 = (0, ∞)^2^. It does not stop mass going off to 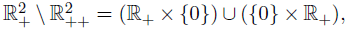 which is exactly what can happen in our case. Thus, their proof only shows the existence of a stationary distribution on 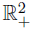 − it does not show the existence of a stationary distribution on 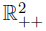. Furthermore, their proof for the uniqueness of a stationary distribution on 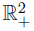 breaks down because their assumption of irreducibility is false. The process (*X*, *Y*) is irreducible on 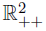, but it is not irreducible on 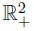 since 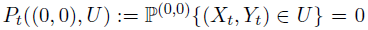 for any open subset *U* that lies in the interior of 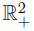. If we work on 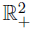, it is not true that the diffusion (*X*, *Y*) has a unique stationary distribution. We can obtain infinitely many stationary distributions on 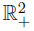 of the form 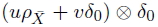 where 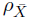 is the unique stationary distribution of 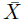 on ℝ_++_ and *u*, *v* ∈ ℝ_+_ satisfy *u* + *v* = 1.

## Appendix D Proof of Theorem 5.1

Assume that the matrix Σ is positive definite and that the dispersion proportion vector *α* is such that *μ* · *α* − *α* · Σ*α*/2 > 0 so that a population playing the strategy *α* persists. Under these assumptions the function *β* ↦ *ℐ*(*α*, *β*) is strictly concave. Hence, by the method of Lagrange multipliers, *ℐ*(*α*, *β*) < 0 for all *β* ≠ *α* and *α*_*i*_ > 0 for all *i* if and only if there exists a constant, which we denote by *λ*, such that

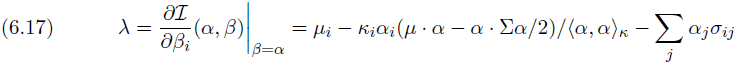

for all *i*. Multiplying (6.17) by *α*_*i*_ and summing with respect to *i*, we get

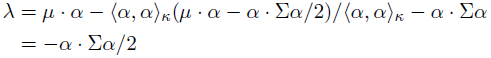

This expression for the Lagrange multiplier and (6.17) provide the characterization of a mixed ESS in equation (5.1) when *α*_*i*_ > 0 for all *i*. The characterization of the more general case of *α*_*i*_ > 0 for at least two patches follows similarly by restricting the method of Lagrange multiples to the appropriate face of the probability simplex.

Suppose that *μ*_*i*_ − *σ*_*ii*_/2 > 0 so that a population remaining in patch *i* and not dispersing to other patches persists. The strategy *α*_*i*_ = 1 and *α*_*j*_ = 0 for all *j* ≠ *i* is an ESS only if

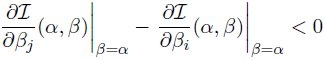

for all *j* ≠ *i*. Evaluating these partial derivatives gives the criterion (5.2) for the pure ESS.

We conclude by considering the case *n* = 2. Define the function *g* : [0, 1] → ℝ by

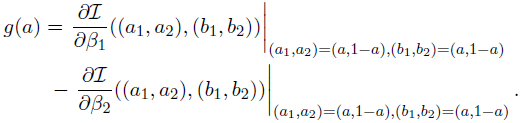

The inequalities (5.2) for the pure strategies (1,0) and (0,1), respectively, correspond to *g*(0) < 0 and *g*(1) > 0, respectively. Hence, when these inequalities are reversed, the intermediate value theorem implies there exists *a* ∈ (0, 1) such that *g*(*a*) = 0. Such an *a* satisfies the mixed ESS criterion (5.1) and, therefore, is an ESS.

## Acknowledgments

The authors thank Dan Crisan, Alison Etheridge, Tom Kurtz, and Gregory Roth for helpful discussions.

## References

J. T. Anderson and M. A. Geber. Demographic source-sink dynamics restrict local adaptation in Elliott’s blueberry (*Vaccinium elliottii*). Evolution, 64:370–384, 2010.

J. P. Beckmann and J. Berger. Using black bears to test ideal-free distribution models experimentally. Journal of Mammalogy, 84:594–606, 2003.

R. S. Cantrell, C. Cosner, D. L. Deangelis, and V. Padron. The ideal free distribution as an evolutionarily stable strategy. Journal of Biological Dynamics, 1:249–271, 2007.

R. S. Cantrell, C. Cosner, and Y. Lou. Evolution of dispersal and the ideal free distribution. Mathematical Biosciences and Engineering, 7:17–36, 2010.

R.S. Cantrell, C. Cosner, and Y. Lou. Evolutionary stability of ideal free dispersal strategies in patchy environments. Journal of Mathematical Biology, 65:943–965, 2012.

P. L. Chesson. General theory of competitive coexistence in spatially-varying environments. Theoretical Population Biology, 58:211–237, 2000.

D.Z. Childs, C. J. E Metcalf, and M. Rees. Evolutionary bet-hedging in the real world: empirical evidence and challenges revealed by plants. Proceedings of the Royal Society B: Biological Sciences, 277:3055–3064, 2010.

C. Cosner. A dynamic model for the ideal-free distribution as a partial differential equation. Theoretical Population Biology, 67:101–108, 2005.

R. Cressman and V. Křivan. Migration dynamics for the ideal free distribution. American Naturalist, 168:384–397, 2006.

R. Cressman and V. Křivan. The ideal free distribution as an evolutionarily stable state in density-dependent population games. Oikos, 119:1231–1242, 2010.

R. Cressman, V. Křivan, and J. Garay. Ideal free distributions, evolutionary games, and population dynamics in multiple-species environments. American Naturalist, 164: 473–489, 2004.

C. P. Doncaster, J. Clobert, B. Doligez, E. Danchin, and L. Gustafsson. Balanced dispersal between spatially varying local populations: an alternative to the sourcesink model. American Naturalist, 150(4):425–445, 1997.

H. Dreisig. Ideal free distributions of nectar foraging bumblebees. Oikos, 72:161–172, 1995.

P. Edelaar and D. I. Bolnick. Non-random gene flow: an underappreciated force in evolution and ecology. Trends in Ecology & Evolution, 27:659 – 665, 2012.

S. N. Ethier and T. G. Kurtz. Markov Processes: Characterization and Convergence. Wiley, Hoboken, NJ, 2005.

S. N. Evans, P. Ralph, S. J. Schreiber, and A. Sen. Stochastic growth rates in spatiotemporal heterogeneous environments. Journal of Mathematical Biology, 66:423–476, 2013.

L. R. Fox and J. Eisenbach. Contrary choices: possible exploitation of enemy-free space by herbivorous insects in cultivated vs. wild crucifers. Oecologia, 89:574–579, 1992.

S. D. Fretwell and H. L. Lucas Jr.. On territorial behavior and other factors influencing habitat distribution in birds. Acta Biotheoretica, 19:16–36, 1969.

A. Friedman. Partial differential equations of parabolic type. Prentice-Hall Inc., Englewood Cliffs, N.J., 1964.

R. Gejji, Y. Lou, D. Munther, and J. Peyton. Evolutionary convergence to ideal free dispersal strategies and coexistence. Bulletin of Mathematical Biology, 74:257–299, 2012.

S. A. H. Geritz, J. A. J. Metz, E. Kisdi, and G. Meszena. Dynamics of adaptation and evolutionary branching. Physical Review Letters, 78:2024–2027, 1997.

J. J. Godin and M. H. A. Keenleyside. Foraging on patchily distributed prey by a cichlid fish (Teleostei, Cichlidae): a test of the ideal free distribution theory. Animal Behaviour, 32:120–131, 1984.

D. G. C. Harper. Competitive foraging in mallards: Ideal free ducks. Animal Behaviour, 30:575–584, 1982.

A. Hastings. Can spatial variation alone lead to selection for dispersal? Theoretical Population Biology, 24:244–251, 1983.

T. O. Haugen, I. J. Winfield, L. A. Vøllestad, J. M. Fletcher, J. B. James, and N. C. Stenseth. The ideal free pike: 50 years of fitness-maximizing dispersal in Windermere. Proceedings of the Royal Society B: Biological Sciences, 273:2917–2924, 2006.

R. D. Holt. On the evolutionary stability of sink populations. Evolutionary Ecology, 11: 723–731, 1997.

R. D. Holt and M. Barfield. On the relationship between the ideal free distribution and the evolution of dispersal. In J. Clobert, E. Danchin, A. Dhondt, and J. Nichols, editors, Dispersal, pages 83–95. Oxford University Press, USA, 2001.

N. Ikeda and S. Watanabe. Stochastic differential equations and diffusion processes, volume 24 of North-Holland Mathematical Library. North-Holland Publishing Co., Amsterdam, second edition, 1989.

J. Jaenike. Genetic and environmental determinants of food preference in Drosophila tripunctata. Evolution, 39:362–369, 1985.

J. Jaenike and R. D. Holt. Genetic variation for habitat preference: evidence and explanations. American Naturalist, 137:S67–S90, 1991.

V. A. A. Jansen and J. Yoshimura. Populations can persist in an environment consisting of sink habitats only. Proceeding of the National Academy of Sciences USA, 95:3696–3698, 1998.

O. Kallenberg. Foundations of Modern Probability. Springer, New York, 2002.

G. S. Katzenberger. Solutions of a stochastic differential equation forced onto a manifold by a large drift. The Annals of Probability, 19:1587–1628, 1991.

V. Křivan. Dynamic ideal free distribution: effects of optimal patch choice on predator-prey dynamics. American Naturalist, 149:164–178, 1997.

J.-F. Le Gall. Applications du temps local aux équations différentielles stochastiques unidimensionnelles. In *Seminar on probability, XVII*, volume 986 of Lecture Notes in Math., pages 15–31. Springer, Berlin, 1983.

X. Li and X. Mao. Population dynamical behavior of non-autonomous Lotka-Volterra competitive system with random perturbation. Discrete and Continuous Dynamical Systems, 24:523–545, 2009.

M. Liu, K. Wang, and Q. Wu. Survival analysis of stochastic competitive models in a polluted environment and stochastic competitive exclusion principle. Bulletin of Mathematical Biology, 73:1969–2012, 2011.

J. Maynard Smith and G. R. Price. The logic of animal conflict. Nature, 246:15–18, 1973.

E. Mayr. Animal species and evolution. Harvard University Press, 1963.

M. A. McPeek and R. D. Holt. The evolution of dispersal in spatially and temporally varying environments. American Naturalist, 6:1010–1027, 1992.

M. Milinski. An evolutionarily stable feeding strategy in sticklebacks. Zeitschrift für Tierpsychologie, 51:36–40, 1979.

T. Oksanen, M. E. Power, and L. Oksanen. Ideal free habitat selection and consumer-resource dynamics. American Naturalist, 146:565–585, 1995.

G. H. Orians and J. F. Wittenberger. Spatial and temporal scales in habitat selection. American Naturalist, 137:S29–S49, 1991.

T. Prout. Sufficient conditions for multiple niche polymorphism. American Naturalist, 102:493–496, 1968.

V. Ravigné, I. Olivieri, and U. Dieckmann. Implications of habitat choice for protected polymorphisms. Evolutionary Ecology Research, 6:125–145, 2004.

H. S. Robinson, R. B. Wielgus, H. S. Cooley, and S. W. Cooley. Sink populations in carnivore management: Cougar demography and immigration in a hunted population. Ecological Applications, 18:1028–1037, 2008.

L. C. G. Rogers and D. Williams. Diffusions, Markov processes, and martingales. Vol. 2. Cambridge Mathematical Library. Cambridge University Press, Cambridge, 2000. Itô calculus, Reprint of the second (1994) edition.

M. L. Rosenzweig. A theory of habitat selection. Ecology, 62:327–335, 1981.

S. J. Schreiber. Evolution of patch selection in stochastic environments. American Naturalist, 180:17–34, 2012.

S. J. Schreiber and M. Vejdani. Handling time promotes the coevolution of aggregation in predator-prey systems. Proceedings of the Royal Society: Biological Sciences, 273: 185–191, 2006.

S. J. Schreiber, L. R. Fox, and W. M. Getz. Coevolution of contrary choices in host-parasitoid systems. American Naturalist, 155:637–648, 2000.

S. J. Schreiber, L. R. Fox, and W. M. Getz. Parasitoid sex allocation affects coevolution of patch selection in host-parasitoid systems. Evolutionary Ecology Research, 4:701–718, 2002.

S. J. Schreiber, M. Benaïm, and K. A. S. Atchadé. Persistence in fluctuating environments. Journal of Mathematical Biology, 62:655–683, 2011.

E. V. Sokurenko, R. Gomulkiewicz, and D. E. Dykhuizen. Source-sink dynamics of virulence evolution. Nature Reviews Microbiology, 4:548–555, 2006.

D. W. Stroock. Partial differential equations for probabilists, volume 112 of Cambridge Studies in Advanced Mathematics. Cambridge University Press, Cambridge, 2008.

R. Tittler, L. Fahrig, and M. A. Villard. Evidence of large-scale source-sink dynamics and long-distance dispersal among Wood Thrush populations. Ecology, 87:3029–3036, 2006.

T. Tregenza. Building on the ideal free distribution. Advances in Ecological Research, 26:253–307, 1995.

M. Turelli, D.W. Schemske, and P. Bierzychudek. Stable two-allele polymorphisms maintained by fluctuating fitnesses and seed banks: protecting the blues in Linanthus parryae. Evolution, 55:1283–1298, 2001.

M. van Baalen and M. W. Sabelis. Coevolution of patch selection strategies of predator and prey and the consequences for ecological stability. American Naturalist, 142: 646–670, 1993.

M. van Baalen, V. Křivan, P. C. J. van Rijn, and M.W. Sabelis. Alternative food, switching predators, and the persistence of predator-prey systems. American Naturalist, 157: 512–524, 2001.

S. Via. Ecological genetics and host adaptation in herbivorous insects: The experimental study of evolution in natural and agricultural systems. Annual Review for Entomology, 35:421–446, 1990.

Z. Zhang and D. Chen. A new criterion on existence and uniqueness of stationary distribution for diffusion processes. Advances in Difference Equations, 2013:13, 2013.

